# Characterizing the Network Architecture of Emotion Regulation Neurodevelopment

**DOI:** 10.1101/773895

**Authors:** João F. Guassi Moreira, Katie A. McLaughlin, Jennifer A. Silvers

## Abstract

The ability to regulate emotions is key to goal attainment and wellbeing. Although much has been discovered about how the human brain develops to support the acquisition of emotion regulation, very little of this work has leveraged information encoded in whole-brain networks. Here we employed a network neuroscience framework to parse the neural underpinnings of emotion regulation skill acquisition while accounting for age in a sample of youth (N = 70, 34 female). Focusing on three key network metrics—network differentiation, modularity, and community structure differences between active regulation and a passive emotional baseline—we found that the control network, the default mode network, and limbic network were each related to emotion regulation ability while controlling for chronological age. Greater network differentiation in the control and limbic networks was related to better emotion regulation ability. With regards to network community structure, more communities and more crosstalk between modules (i.e., less modularity) in the control network were associated with better regulatory ability. By contrast, less crosstalk (i.e., more modularity) between modules in the default mode network was associated with better regulatory ability. Together, these findings highlight possible whole-brain connectome features that support the acquisition of emotion regulation in youth.

## Introduction

Emotions promote adaptive behavior but can become detrimental if left unchecked (Gross, 2015), underscoring the importance of emotion regulation. Emotion regulation acquisition is a protracted process that emerges across development, and is highly consequential for wellbeing (Calkins, 1994; Casey et al., 2008; Thompson et al., 2008). Over the past fifteen years, substantial efforts have been made to examine brain regions involved in the development of emotion regulation, with most work focusing on activation within, or dynamics between, the amygdala, ventromedial prefrontal cortex (vmPFC), and lateral prefrontal cortex (lPFC) (Ernst, Pine, & Hardin, 2006; Gee et al., 2014; Kim et al., 2013; McRae et al., 2012; Perino, Miernicki, & Telzer, 2016; Pitskel, Bolling, Kaiser, Crowley, & Pelphrey, 2011; Silvers et al., 2017). However, surprisingly little is known about the extent to which whole-brain networks—suites of brain regions that reliably activate in concert (e.g., default mode, control)—support the acquisition of emotion regulation skills. This omission is noteworthy given work in adults showing that many emotional, as well as non-emotional, processes and behaviors are supported by whole-brain networks (Gratton et al., 2018; Kleckner et al., 2017). Leveraging network neuroscience may help to fill at least two major gaps in our knowledge of emotion regulation neurodevelopment. First, this approach may improve our understanding of how correlated but distinct developmental phenomena such as chronological age and emotion regulation ability are represented in the brain. Second, neuroscientific theories of emotion regulation implicitly posit dynamic interactions between brain regions during active emotion regulation (Ochsner, Silvers, & Buhle, 2012), yet only recently have imaging studies of emotion regulation begun to explicitly test this assumption (Zhang, Padmanabhan, Gross, & Menon, 2019). Doing so from a whole-brain network perspective will provide more specific and precise evidence to be used in evaluating said theories.

Network neuroscience is a promising approach for parsing distinct but related developmental phenomena (Richardson et al., 2018). The current study focused on two network properties that have been particularly important for understanding development in other domains and thus may be fruitful for pinpointing the neurodevelopment of emotion regulation ability. The first property is network differentiation—the extent to which activity within a network is similar or distinct across task states, perhaps reflecting a network’s “specialization” for a given task. Examining network differentiation in combination with behavioral assessments of emotion regulation can help determine which networks become specialized to support skill development (i.e., improved emotion regulation ability). While some patterns of network differentiation may correspond with changes in specific skills, others that support more general aspects of development may show increased specialization with age rather than tracking with performance on a given task. For instance, prior research shows greater specialization among pain perception and mentalizing networks tracks with chronological age, but not skills related to representing others’ states (Richardson et al., 2018). Extrapolating to emotion regulation, this implies that some networks might support domain-general development with chronological age (e.g., the default mode network), whereas others specifically support emotion regulation abilities (e.g., the salience network). The second property is network community structure—or, how brain networks are divided into sub-structures (“communities”, “modules”; Sporns, 2010; Yeo et al., 2011). In the context of emotion regulation, a network’s modules may support different emotion regulation sub-processes (e.g., working memory). We assessed network community structure here by examining the number of communities that emerged in a given network between task states as well as modularity, the strength of intra-versus inter-module connections. A network’s community structure (number of communities, modularity) and the suite of psychological processes it supports, might track differentially with age and ability, helping reveal the manner in which domain-general maturation (chronological age) differs from growing expertise in a skill (ability). Given that other lines of research have successfully leveraged network differentiation and community structure metrics to build increasingly accurate models of brain function, cognitive skills, and development (Baum et al., 2017; Richardson et al., 2018; Sizemore et al., 2018), it stands to reason that such an approach may also benefit the study of emotion regulation development.

In the present study, we used the three aforementioned metrics of network activity (network differentiation, number of communities, modularity) in conjunction with ridge regression to predict regulation ability in a sample of youth who completed an emotion regulation task during functional magnetic resonance imaging (fMRI) scanning. Because our questions were exploratory in nature, our study was guided by open questions about how network properties relate to emotion regulation ability beyond chronological age. However, given prior neuroimaging work implicating prefrontal-subcortical circuitry in emotion regulation development (Casey, 2015; Silvers et al., 2017), we generally expected that limbic and prefrontal control networks were particularly likely to explain differences in regulatory ability.

## >Methods

### Analytic Overview

In the current study, we scanned 70 children and adolescents with fMRI while they completed four runs of a cognitive reappraisal task that involved alternately regulating (regulate condition) and reacting naturally (emotional baseline condition) to a series of negative and neutral images. Cognitive reappraisal—reframing a stimulus to alter its emotional impact—was our chosen form of emotion regulation study because it is widely-studied, is highly consequential for adjustment (e.g., Troy et al., 2010), its abilities are known to improve with age during childhood and adolescence (Gross, 1998; McRae et al., 2012; Silvers et al., 2012).

The following procedure was conducted to derive the network metrics we would use for both aims articulated above. Following visual and statistical assessment of data quality and preprocessing, we prepared the data for analysis by extracting beta-series from each node (i.e., ROI) across all networks defined by the Schaeffer 17 parcellation scheme (more information on parcellation selection below). Various metrics from each network were next calculated with the goal of using said metrics to build classifiers that could predict age and emotion regulation ability Correlations among each networks’ beta-series were performed and used, in conjunction with representational similarity analysis (RSA) and graph theory, to yield our metrics of network activity. These metrics included network differentiation between task states (regulate v. emotional baseline; RSA based) and two measures of network community structure: 1) differences in the *number of modules* within a network between task states (regulate v. emotional baseline), and 2) differences in *modularity* between task states (the degree of ‘crosstalk’ between modules for regulate v. emotional baseline). Notably, the number of modules and modularity are related but distinct—the former tells us how many ‘divisions of labor’ occur in a network and the latter reflects how rigidly the network appears to follow such divisions. We reiterate here that our inclusion of, and comparison to, a control conditional (a passive emotional baseline) is critical for making inferences about emotion regulation rather than broader emotional processes.

Afterwards we submitted connectivity-based metrics from the top specifications to ridge regression analyses predicting age and emotion regulation ability. Modeling decisions comprising parcellation, edge defining threshold, and whether to detrend beta-series were decided by running a ridge regression model on each specification and selecting the specification that yielded the lowest RMSE. After selecting the appropriate specification, we performed inference over parameter estimates by building bootstrapped confidence intervals. Further details follow.

### Participants

Participants for the current study were drawn from a larger sample of individuals enrolled in a longitudinal neuroimaging study about childhood maltreatment and affective neurodevelopment. Youth in the current analyses were selected from a non-maltreated community control group who were screened with the following exclusion criteria: childhood maltreatment/violence, presence of a developmental disorder (e.g., autism), psychotropic medication use, and IQ < 75. To qualify for inclusion in the current report, participants were required to have anatomical images free of abnormalities or artifacts, low levels of motion during the scan (addressed below), and accompanying behavioral data from our scanning emotion regulation paradigm. Our final sample included 70 youth (34 female) ranging in age from 8.08 to 17.00 years (Mean age = 12.7). The study was conducted in a large, metropolitan area in the Pacific Northwest region of the United States. Youth and their parents provided written assent and consent, respectively, in accordance with the University of Washington’s Institutional Review Board. Participants were remunerated $75 after completion of the scan. Data and analysis code can be accessed on the Open Science Framework (osf.io/bfg3d).

#### Experimental Design

##### fMRI Emotion Regulation Paradigm

Participants completed a variation on a common and widely used emotion regulation task for youth while undergoing fMRI (Silvers, Shu, Hubbard, Weber, & Ochsner, 2015). Participants passively viewed or effortfully regulated emotional responses to a series of developmentally-appropriate negative and neutral images (modeled after the International Affective Picture System (IAPS); Lang, Bradley, & Cuthbert, 2008). Each trial began with an instructional cue (2s duration), ‘Look’ or ‘Far’. The former instructs participants to passively view a subsequent stimulus whereas the latter prompts participants to regulate it via a psychological distancing variant of cognitive reappraisal. Following the cue, participants viewed a neutral or aversive stimulus. Aversive stimuli were either paired with ‘Look’ or ‘Far’. Neutral images were always paired with the ‘Look’ cue; these images are not of interest to the current study and will not be discussed further. Stimuli were displayed for 6-10s (jittered). A 5-point Likert scale was presented thereafter to collect participants affective responses (4s). Lastly, a jittered ITI concluded each trial (1.5-6.5s). Prior to scanning, participants were trained extensively by experimenters to ensure they understood the task and were comfortable completing it, and afterwards completed a practice run. Stimuli were purchased from a stock photography website (https://shutterstock.com) and were normalized in a sample of youth (120, ages 6-16, 50% female) based on valence, arousal, and dominance. Stimuli and normed ratings are available online (osf.io/43hfq/). The most negatively rated images were selected for use in the current study. Participants completed four runs, each lasting approximately 220s (rungs ranged between 110-115 4D volumes). Each task condition (‘Look’ – Negative; ‘Far’ – Negative; ‘Look’ – Neutral) was presented 20 times over the four runs (5 per run). The task was programmed and presented in E-Prime (Psychology Software Tools, Inc., http://www.pstnet.com).

##### fMRI Data Acquisition

If participants were younger than 12 years of age or exhibited any nervousness about scanning, they were taken to a mock MRI scanner in order to familiarize with them with scanning environment and to be trained on how to minimize head motion in the scanner. Additionally, in preparation for the scan, participants were packed into the head coil with an inflated, head-stabilizing pillow to restrict movement.

Images were acquired at the University of Washington’s (Seattle) Integrated Brain Imaging Center on a 3T Phillips Achieva scanner. A 32-channel head coil was used in conjunction with parallel image acquisition. A T1-weighted, magnetization-prepared rapid acquisition gradient echo (MPRAGE) image was acquired for registration purposes (TR=2530ms, TE=1640-7040μs, 7° flip angle, 256 mm^2^ FOV, 176 slices, 1 mm^3^ isotropic voxels). Blood oxygenation level dependent (BOLD) signal during functional runs was recorded using a gradient-echo T2*-weighted echoplanar imaging (EPI) sequence. Thirty-two 3-mm thick slices were acquired parallel to the AC-PC line (TR=2000ms, TE=30ms, 90° flip angle, 256 x 256 FOV, 64 x 64 matrix size).

##### fMRI Data Pre-Processing

The FMRIB Software Library package (FSL, version 5.0.9; fsl.fmrib.ox.ac.uk) was used for pre-processing and later analysis. Prior to pre-processing, data were visually inspected for artifacts, anatomical abnormalities, and excessive head motion. Pre-processing began by using the brain extraction tool (BET) to remove non-brain tissue from images and estimating the extent of participant head motion by using FSL Motion Outliers to record volumes that exceed a 0.9 mm framewise displacement (FD) threshold (Siegel et al., 2014). Fortunately, head motion was minimal: The average number of volumes exceeding our FD threshold per run, per participant was 3.014 (*SD* = 6.09, range = 0 - 30.75). Head motion as not appreciably correlated with age or emotion regulation ability scores (both *rs* < |.05|). Afterwards, we corrected for head motion by spatially realigning volumes using MCFLIRT and then hi-pass filtered the data (100s cutoff). We pre-whitened the data to correct for temporally autocorrelated residuals.

##### Network Definition

We initially began with two prospective network parcellations to choose from: the Schaefer 7 network parcellation (100 ROIs) and the Schaefer 17 network parcellation (400 ROIs) (Schaefer et al., 2018). For both parcellations, spheres (4 mm radius) were drawn across all peaks in order to extract data necessary to compute connectivity (described in the next section). A 4mm radius was used to minimize overlap between spheres. We made the decision to extract data from identically sized spheres to ensure that differential effects between networks were not driven by differences in volume. We ultimately selected the Schaefer 17 parcellation based on evaluating fit statistics from several different possible models (see ‘Statistical Analysis’ section).

In addition to the networks in the Schaefer7 and Schaefer17 parcellations, we also included ROIs from a ‘cognitive reappraisal network’ (CRN; Buhle et al., 2014). These ROIs are commonly observed when individuals engage in cognitive reappraisal, the emotion regulation strategy employed here. We decided to include the CRN because it is the most relevant network to the emotion regulation task incorporated in the present study. Recent work in both adult and developmental samples suggests that task-specific information—and even activity from task-specific networks—is privileged over domain-general information and thus may be informative when differentiating between task ability and chronological age (Brody et al., 2019; Greene, Gao, Scheinost, & Constable, 2018). Nodes in the CRN were identified using the same procedure described in Guassi Moreira, McLaughlin, & Silvers, 2019.

#### Estimating Network Connectivity Metrics

##### Network Connectivity via Beta-Series

In order to obtain metrics of network-level activity, we first modeled the task data within-subjects using a Least Squares Single (Mumford et al., 2014) design to yield a beta-series (Rissman et al., 2004). For each run within an individual subject, a fixed-effects GLM was created for each ‘Far’ (regulation of aversive stimuli) and ‘Look-Negative’ (passive observation of aversive stimuli) ‘target trial’. Thus, a separate GLM was created for each individual target trial. Each GLM consisted of a single-trial regressor which modeled that GLM’s target trial, a nuisance regressor modeling all other events in the target trial’s condition, regressors for the other two conditions—each modeled as they normally would be in a standard univariate analysis. For example, the first ‘Far’ trial for a participant would receive its own GLM wherein it was modeled in a single regressor; all other ‘Far’ trials were placed in a separate nuisance regressor while ‘Look-Negative’ and ‘Look-Neutral’ trials modeled in their own respective regressors. Afterwards, a new GLM was created for the *second* ‘Far’ trial, where it was the ‘target trial’ and modeled in a single-event regressor and the *first* trial is now modeled in the separate nuisance regressor with other trials of that type. FSL’s extended motion parameters estimated from MCFLIRT (rotation+translation, derivatives, and squares of each) and additional volumes that exceeded our FD = 0.9 mm threshold were also added as regressors of non-interest to each first-level LSS GLM.

Following estimation of the LSS GLMs, we used parameter estimates from each trial-specific GLM to create linear contrast images comparing both trial types of interest (‘Far’ & ‘Look-Negative’), respectively, to baseline. We then extracted estimates of activation for both trial types across a series of ROIs nested within six different networks (see below). This yielded two *t* x *p* x *n* arrays (one for each task condition), where entries to each cell represent the average parameter estimate at the *t*-th trial for the *p*-th ROI in the *n*-th network. For every network, beta-series amongst all its ROIs were correlated amongst each other (Spearman’s Rho), yielding two connectivity matrices (one for each trial type) per network. These matrices describe each network’s connectivity for a given task condition (i.e., connectivity matrix), and we refer to them as network connectivity profiles (See Figure 1 for an overview).

**Figure 1.**
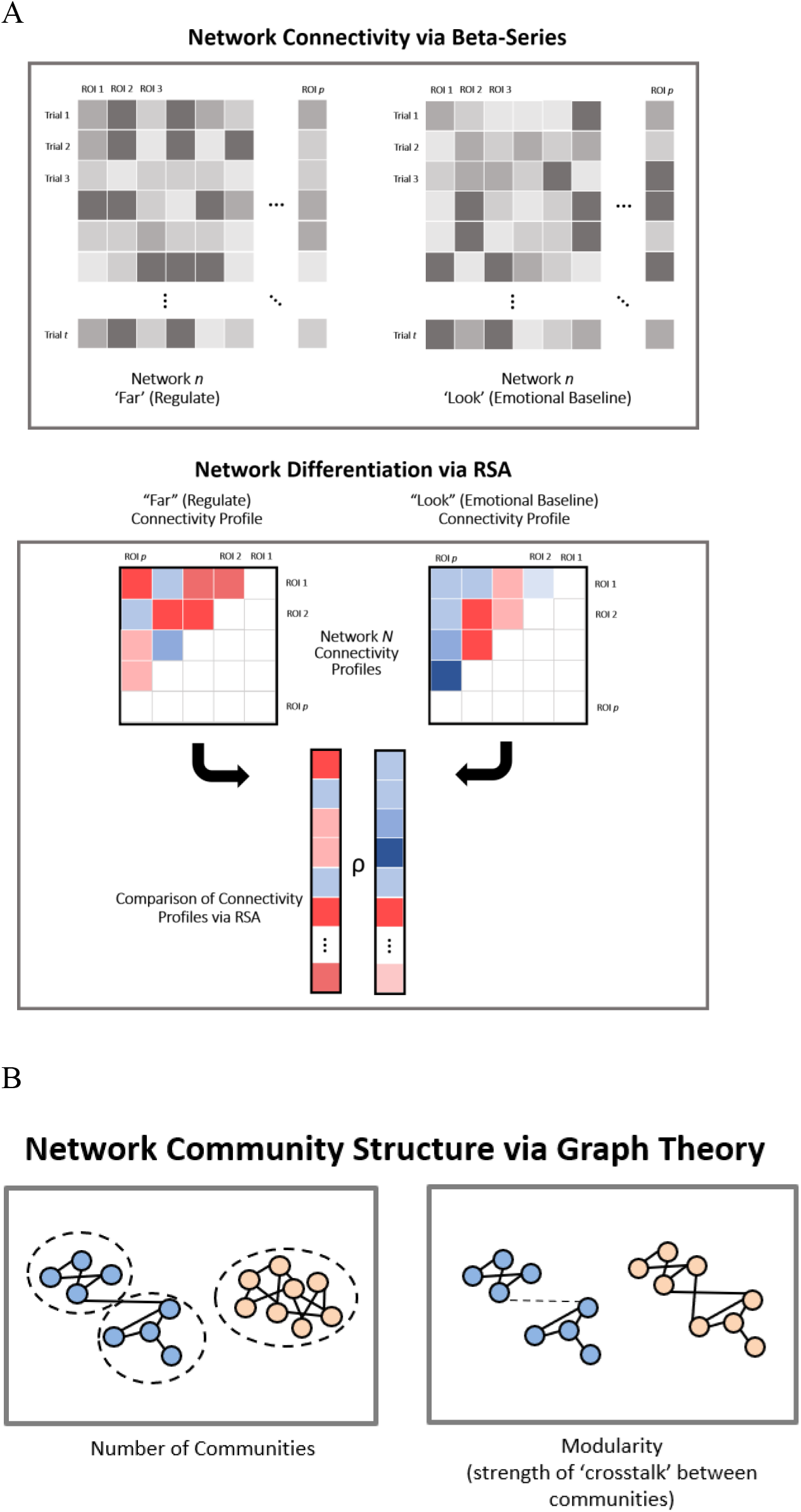
Overview of network activity metrics estimation. *Note*. **A.** Top panel (Network Connectivity via Beta-Series) depicts the beta-series estimated for each task condition (regulate (‘Far’), emotional baseline (‘Look-Negative’)), for the *n-th* network. Each column corresponds to the *p*-th ROI, each row to the *t*-th trial, and each entry is the average beta-value for a given ROI at a given trial. Beta-series for each subject served as the substrates for estimating our network activity metrics of interest, depicted in the bottom row of panels. The middle panel (Network Differentiation via RSA) depicts process of computing network differentiation. The two boxes represent connectivity profiles (matrices) for each condition for the *n*th network. Each entry to the matrix is the connectivity estimate between a given pair of ROIs. Unique off-diagonal elements were vectorized and then correlated (Spearman’s Rho) to yield a single, scalar value of network connectivity similarity. The bottom panel (Network Community Structure via Graph Theory) depicts graphs for each task condition for the *n*-th network; each node represents an ROI within a given network, whereas each edge is the connectivity between a pair of ROIs. This step allowed us to compute the number of network communities (i.e., number of modules/communities in each network) and modularity. **B.** The left panel shows two illustrative diagrams of communities, or neighborhoods of highly interconnected nodes. The network with blue nodes has two communities while the network with yellow nodes has just one. The right panel illustrates two networks that differ in terms of modularity, or the strength of cross-talk between different communities, represented by edges between communities. The network with blue nodes has greater modularity or less cross-talk because the connection between the two communities is weak (indicated by the dashed line), whereas the network with yellow nodes has less modularity or more cross-talk because the connection between the two communities is strong (indicated by two connections between neighborhoods, solid lines). Network connectivity matrices were thresholded to create adjacency matrices and subsequently graphs. Estimates of community structure and modularity were compared across both task conditions for each network.

#### Network Differentiation

##### Using Representational Similarity Analysis to Quantify Network Differentiation

The first feature of network activity we chose to examine was differentiation. In order to estimate network differentiation, we compared profiles of network connectivity when youth were engaging in emotion regulation versus when they were simply viewing emotional stimuli using representational similarity analysis (RSA) on our two task-specific connectivity matrices (Kriegeskorte, 2008; Kriegeskorte et al., 2008). Traditionally, RSA is a form of multi-voxel pattern analysis that relies on the presumption that multi-voxel activation patterns reliably contain information about a specific stimulus (Etzel et al., 2013; Kriegeskorte et al., 2009). Comparison of two stimulus patterns (via correlation) in RSA can reveal the extent to which representations of stimuli encode unique or similar information. However, one is not bound to use RSA solely with multivariate patterns. In fact, RSA can be used to compare any two different representations because information about pattern representations is ultimately abstracted away from the modality in which it was initially acquired and transformed forced into a common space. To this point, prior research has used RSA to compare voxelwise patterns in the brain to behavioral patterns (Parkinson et al., 2017), behavioral patterns to other behavioral patterns (Brooks & Freeman, 2018; Stolier et al., 2018), and, most importantly for our purposes, representations of brain connectivity patterns (Lee et al., 2017). We built upon and extended these prior implementations of RSA for our purposes in comparing network connectivity states between task conditions.

For each network and task condition, a *r* x *r* connectivity matrix was constructed, where *r* is the number of ROIs in the network. Representational similarity analyses involved vectorizing the off-diagonal elements of each task condition’s various connectivity matrices, Fisher transforming them (for variance stabilization), and then correlating the elements (Spearman). This yielded a scalar value that describes the extent to which a given network’s connectivity profile differs between task conditions (see Figure 1). An extreme high value indicates that the pattern of connectivity across the entire network remains consistent during ‘Far’ (Regulate) and ‘Look-Negative’ (Emotional Baseline) trials; An extreme low value (anticorrelation) indicates a differentiated connectivity profile between the two task states.

##### Network Community Structure

The next feature of network activity we examined was community structure between task conditions (e.g., when regulating versus when passively viewing emotional stimuli) (Sporns, 2010). That is, we examined the number of communities within a network, or subclusters comprised of densely interconnected nodes within the network, as well as crosstalk (i.e., modularity) between communities.

##### Community Number Differences Between Network States

When examining community structure, we first quantified the number of communities within each network as a function of task condition. To achieve this, we used the aforementioned connectivity matrices to create undirected, weighted adjacency matrices. An adjacency matrix summarizes a graph—each row and column represent the ROIs (nodes) that comprise the network, while an entry into the off-diagonal elements of the matrix denotes the presence or absence of a connection (edge) between two nodes. It is customary to set an *edge defining threshold*, a value at which all correlation estimates (e.g., correlation between two ROIs’ beta-series) at or falling below the threshold are set to zero, and all estimates above the threshold are set to 1 (unweighted edges) or left at their original value (unweighted edges) (Bolt, Nomi, Yeo, & Uddin, 2017). In our case, we used weighted edges. We varied the edge defining threshold as part of our model specification selection procedure (see Statistical Analysis section).

We then estimated the number of communities in each task condition, for all networks, for all subjects using the R igraph package’s walktrap clustering algorithm (Csardi & Nepusz, 2006; Pons & Latapy, 2005) in R (R Core Team, 2014). The walktrap clustering algorithm (Pons & Latapy, 2005) estimates community structure in a graph using random walks—step-wise paces from one node to another on a graph according to random chance. The algorithm operates on the principle that the probability of taking a walk (i.e., moving over) from one node to another depends on the number of shared connections between nodes. This ultimately means that walks are more likely to become ‘trapped’ inside a community than a non-community due to the dense interconnections between nodes that define community membership. The number of a given network’s communities across both task conditions were extracted from the graphs and differenced (‘Far’ – ‘Look-Negative’), resulting in six metrics of community structure, one per network, per participant. A higher value on this metric indicates a greater number of observed communities within a network during emotion regulation, compared to passively viewing emotional stimuli.

##### Modularity Differences Between Network States

Our final network metric was the difference in modularity between task conditions for a given network. Building upon the concept of community structure, a graph is said to be highly modular if it exhibits relatively dense interconnections within its communities and relatively sparse connections between them. On the other hand, a graph that is low in modularity has nodes that have relatively equal connections within and between their communities. In other words, modularity can be thought to measure the extent to which “cross-talk” exists between the communities within a network (i.e., a ratio between intra-versus inter-community ties). Modularity metrics across both task states in each network across all participants were obtained using the communities identified in the prior section. The same differencing was taken (‘Far’ – ‘Look-Negative’) to obtain a metric of modularity differences between task conditions across each participant’s connectome. Notably, because our focus was on the degree to which information about development and emotion regulation ability is encoded across a set of brain networks, we opted to focus on *within-network* modularity, rather than *between*-network modularity.

#### Statistical Analysis

##### Selecting a Parcellation and Other Modeling Decisions

We were faced with a number of different modeling decisions, including choosing a parcellation (Schaefer7 or Schaefer 17), an edge-defining threshold (ρ = 0.4, 0.5, 0.6), and whether to linearly detrend the beta-series via OLS regression (detrend, do not detrend). In order to do this, we estimated a model using all possible specifications (12 total) computed their associated root mean square error (RMSE) as a metric of prediction accuracy. The final model was estimated using specifications that yielded the lowest RMSE. Notably, the RMSE was weighted by degrees of freedom to avoid bias towards model with more predictors. Both the Schaefer7 and Schaefer17 networks included CRN ROIs.

##### Model Building Using Ridge Regression

We used ridge regression to estimate associations between network activity metrics and emotion regulation ability (controlling for age). Ridge regression was selected because (i) we had continuous dependent variables, and because ridge regression (ii) effectively estimates parameters in a model with many predictors, (iii) handles highly correlated predictor variables, and (iv) most importantly, is less susceptible to overfitting and therefore has better out of sample predictive accuracy (McNeish, 2015; Murphy, 2012). Ridge regression differs from traditional OLS regression in that its loss function contains an extra penalty term (Penalty *b_Ridge_* = λΣ*b_j_*^2^). This has the effect of biasing, or shrinking, coefficients towards zero, especially those with inappropriately large magnitudes that drive overfitting. This added bias, introduced via what is known as *l2 regularization*, helps prevent overfitting by decreasing sample to sample variability in regression coefficients. This has the effect of producing more generalizable models while satisfying the traditional aim of helping make inferences about population parameters due to enhanced certainty in parameter estimates (i.e., narrower sampling distribution of coefficient estimates).

Here we implemented ridge regression using the `glmnet()` function in R. Notably, ridge regression requires a tuning parameter (λ) that controls the degree of regularization. In our case, we used the `cv.glmnet()` function, which finds the optimal λ value via cross-validation. To facilitate statistical inference, we computed 95% bootstrapped confidence intervals (percentile) around parameter estimates (5,000 bootstrapped iterations).

## Results

### Behavioral Results

Behavioral results from our emotion regulation task have been reported elsewhere (see Guassi Moreira et al., 2018), but we briefly recapitulate them here for convenience. Participants’ average negative affective ratings during the ‘Look-Negative’ and ‘Far’ trials were 2.48 (SD = .562) and 1.95 (SD = .499), respectively. This difference was statistically significant (*t*(69) = 10.37, *p* < .001). Participants’ ability to engage in emotion regulation, quantified as the percent reduction in negative affect for the regulate condition relative to the emotional baseline condition, improved with age (*r* = 0.344, *p*<.01).

#### Using network approaches to parse chronological age and emotion regulation ability

##### Model Selection

The specified model with the lowest RMSE used the Schaefer 17 parcellation, an edge defining threshold of ρ = 0.4, and linearly detrended beta-series (via OLS). This model’s RMSE was decidedly lower than those of the next lowest models (0.808 vs 0.916+), suggesting superior model fit.

##### Predicting Emotion Regulation Ability

Ridge regression parameters and bootstrapped confidence intervals are listed in Table 1. When predicting emotion regulation ability from our network connectivity metrics, we found that network differentiation in Control Network (subnetwork C) and Limbic Network (subnetwork B) were each predictive of emotion regulation ability such that greater differentiation between these networks’ connectivity profile during active regulation compared to emotional baseline was related to better emotion regulation ability. Differences in network modularity during emotion regulation compared to emotional baseline were also related to emotion regulation ability. Specifically, greater modularity in Default Mode Network (subnetwork B) was associated with increased emotion regulation ability whereas the opposite was true for the Control Network (subnetwork C). Only community differences in Control Network (subnetwork B) were related to emotion regulation ability, such that more communities during regulation versus emotional baseline were associated with better emotion regulation ability. Notably, all of these effects were significant while statistically adjusting for all terms in the model, including age. Age was directly associated with emotion regulation ability after statistically adjusting for all network terms.

**Table 1.** Ridge regression results predicting emotion regulation ability

| Term | Parameter Estimate | 95% CI | Term | Parameter Estimate | 95% CI |
| --- | --- | --- | --- | --- | --- |
| simCoNA | -0.466 | [-0.179, 0.079] | modSaVANA | 1.088 | [-0.015, 0.239] |
| simCoNB | 0.345 | [-0.058, 0.195] | modSaVANB | 0.003 | [-0.127, 0.105] |
| **simCoNC** | **-1.646** | **[-0.341, -0.036]** | modSMNA | 0.856 | [-0.042, 0.186] |
| simDMNA | 0.215 | [-0.089, 0.14] | modSMNB | -0.121 | [-0.151, 0.1] |
| simDMNB | -0.118 | [-0.13, 0.144] | modTPN | 0.462 | [-0.069, 0.176] |
| simDMNC | 0.835 | [-0.007, 0.254] | modVCN | 0.111 | [-0.096, 0.126] |
| simDANA | 0.117 | [-0.112, 0.126] | modVPN | -0.208 | [-0.118, 0.109] |
| simDANB | -1.078 | [-0.245, 0.015] | modCRN | 0.389 | [-0.073, 0.172] |
| simLiNA | -0.174 | [-0.126, 0.117] | comCoNA | 0.481 | [-0.11, 0.179] |
| **simLiNB** | **-1.354** | **[-0.256, -0.022]** | **comCoNB** | **1.528** | **[0.017, 0.239]** |
| simSaVANA | -0.195 | [-0.14, 0.097] | comCoNC | 0.443 | [-0.114, 0.144] |
| simSaVANB | 0.041 | [-0.111, 0.12] | comDMNA | 0.445 | [-0.081, 0.192] |
| simSMNA | -0.284 | [-0.135, 0.105] | comDMNB | -0.110 | [-0.156, 0.116] |
| simSMNB | 0.220 | [-0.093, 0.153] | comDMNC | -0.903 | [-0.252, 0.038] |
| simTPN | -0.762 | [-0.211, 0.051] | comDANA | 0.135 | [-0.101, 0.159] |
| simVCN | -0.014 | [-0.137, 0.129] | comDANB | 0.805 | [-0.111, 0.177] |
| simVPN | 0.330 | [-0.096, 0.127] | comLiNA | 0.365 | [-0.083, 0.188] |
| simCRN | -0.734 | [-0.222, 0.06] | comLiNB | 0.372 | [-0.087, 0.173] |
| modCoNA | -0.739 | [-0.179, 0.055] | comSaVANA | 0.564 | [-0.104, 0.148] |
| modCoNB | 0.312 | [-0.115, 0.131] | comSaVANB | 0.851 | [-0.05, 0.211] |
| **modCoNC** | **-1.597** | **[-0.333, -0.018]** | comSMNA | -0.021 | [-0.173, 0.084] |
| modDMNA | 0.462 | [-0.075, 0.143] | comSMNB | 0.062 | [-0.167, 0.119] |
| **modDMNB** | **1.971** | **[0.055, 0.305]** | comTPN | 0.748 | [-0.039, 0.227] |
| modDMNC | -0.429 | [-0.157, 0.071] | comVCN | -0.914 | [-0.227, 0.026] |
| modDANA | 0.381 | [-0.066, 0.151] | comVPN | 0.868 | [-0.027, 0.197] |
| modDANB | 0.163 | [-0.112, 0.163] | comCRN | 0.465 | [-0.076, 0.197] |
| modLiNA | -0.559 | [-0.193, 0.087] | AGE | 1.922 | [0.042, 0.301] |
| modLiNB | 0.310 | [-0.115, 0.12] | - | - | - |

*Note.* ‘sim’ refers to network differentiation (obtained via RSA); ‘mod’ refers to differences in modularity (regulation – baseline); ‘com’ refers to differences in community structure, i.e., the number of communities (regulation – baseline). Greater differentiation is indicated by relatively lower similarity scores (i.e., higher similarity indicates that a network’s connectivity is less differentiated between task states). CoN refers to control network; DMN refers to default mode network; DAN refers to dorsal attention network; LiN refers to limbic network; SaVAN refers to salience/ventral attention network; SMN refers to somatomotor network; TPN refers to temporal-parietal network; VCN refers to visual center network; VPN refers to visual peripheral network; CRN refers to cognitive reappraisal network. Letters next to each network indicate subnetwork. 95% confidence intervals reflect bootstrapped percentile intervals (5,000 bootstraps). Predictors whose 95% CI did not include zero, thus indicating statistical significance, are bolded.

Given that multiple subnetworks of Control Network showed significant associations with emotion regulation ability, we decided to examine the correlation between Control Network subnetwork predictors. While the use of ridge regression theoretically obviates concerns about collinearity among predictors, it does not address the degree to which these subcomponents represent distinct entities. Fortunately, none of the Control Network subnetworks were highly correlated across the three metrics (differentiation *r*s: .26 - .40; modularity *r*s: -.07 - .26; community *r*s: .11 - .38), indicating that these subnetworks are related but distinct units, in the context of emotion regulation.

Last, in order to determine the variance in ability that was uniquely predicted by the aforementioned network metrics as well as age, we calculated squared semi-partial correlations. Network metrics each uniquely accounted for between 4.49% and 9.84% of the variance in emotion regulation ability, while age uniquely accounted for 4.65% of the variance in emotion regulation ability (Table 2). Because these are squared semi-partial correlations, these estimates of variance accounted for reflect statistical adjustment of the other significant terms in the model.

**Table 2.** Semi-partial correlations between select brain network metrics and emotion regulation ability

| Predictor | Squared Semi-Partial Correlation |
| --- | --- |
| Age | 4.65% |
| Control Network differentiation | 7.18% |
| Limbic Network differentiation | 4.64% |
| Control Network modularity | 4.49% |
| Default Mode Network modularity | 9.84% |
| Control Network community number | 5.37% |

*Note.* Semi-partial correlations reflect unique variance that each predictor in the table explains in emotion regulation ability after partialling out shared variance with the other predictors. In other words, they are Pearson correlations between emotion regulation ability and the residuals of a given predictor by the others. They are reported here in a percent metric to facilitate interpretation of unique variance accounted for by each predictor.

## Discussion

We set out to interrogate the neurodevelopment of emotion regulation using network neuroscience by parsing brain network features that contribute to correlated but distinct developmental processes (chronological age and acquisition of emotion regulation abilities). This work builds upon past network neuroscience studies of development to better understand task-specific aspects of maturation (Rudolph et al., 2017). We found that emotion regulation ability showed significant associations with whole-brain network metrics in the control network, default mode network, and the limbic network. Generally, because greater network differentiation and modularity in these networks was associated with better regulation ability, it suggests these networks develop specialized features that lend themselves to better implementing emotion regulation. Importantly, these combined network properties explained a substantial amount of the variance in emotion regulation ability over and above age. These findings have implications for our basic understanding of the neurodevelopment of emotion regulation, and how brain networks may broadly change and interact across cognitive and emotional development.

In the present study, we were able to explain variance in emotion regulation ability using network activity metrics (cumulative variance explained by brain predictors after controlling for age = 36.17%), even after accounting for age (variance explained after controlling for brain predictors = 4.65%). These results suggest that task-based network activity encodes information about skill development associated with a given task rather than domain general features of development (i.e., age). We found that greater differentiation in the Control and Limbic Networks were associated with increased emotion regulation ability. These results are directly in line with what is posited by dominant neurobiological models of cognitive emotion regulation functions (Etkin, Büchel, & Gross, 2015; Ochsner et al., 2012). These models posit that bottom-up affective signals generated in limbic regions are modulated by top-down cognitive processes in prefrontal cortical areas. Our data support this notion by showing that youths with better regulation skills show more differentiated patterns of connectivity in the Control and Limbic Networks. That is, connectivity in these networks looks different when regulating than when passively experiencing emotions.

We also found that differences in Default Mode and Control Network modularity were associated with regulation ability, albeit in different ways. Greater modularity in the Default Mode Network during regulation, relative to baseline, was related to better ability, whereas the opposite trend was found with Control Network. This is consistent with recent neurodevelopmental findings showing that increasing functional subdivision of the DMN facilitated better mentalizing abilities on a theory of mind task (Richardson et al., 2018), as well as clinical work linking enhanced cross-network connectivity and activity in the Default Mode Network with emotion regulation disorders such as Major Depression (Liu et al., 2012; Sheline et al., 2009). Because cognitive reappraisal relies on higher order executive functions such as self-referential thought that are widely considered to be supported by DMN (Buckner & Carroll, 2007), it is possible that the anatomical substrates of the network segregate to support functional specialization. While the opposite trend with the Control Network can appear initially contradictory, it is worth noting that evincing a greater number of network communities in Control Network (relative to baseline) were related with better regulatory ability. It is noteworthy that prior work found that increasing structural modularity of the Control Network is associated with increased cognitive skills across age (Baum et al., 2017). In tandem, these results suggest that greater functional subdivision in the Control Network (more communities) *is* related to better regulatory ability, but only when functional modules are engaging in enough ‘cross-talk’ (less modularity). Thus, the Control Network best supports developing emotion regulation abilities when it is characterized by ready communication between specialized modules. Similar to the DMN findings, the specific subnetwork of the Control Network involved most consistently in our study was comprised of individual ROIs that have been implicated in visuospatial imagery and self-referential processing (Cavanna & Trimble, 2006). These are two psychological processes that are heavily involved in the distancing variant of reappraisal we employed and further suggest functional specialization supports fine-tuning of the cognitive skills needed to engage in reappraisal.

Overall, our findings speak to the nature of neurodevelopment. Specifically, they suggest that maturation is driven by changes in network states. Rather than viewing neural network development as monolithic, this perspective suggests that different psychological states – for example, regulating emotions versus not regulating emotions – elicit unique and discrete configurations of activity and connectivity within a single network (Shine, et al., 2019a b)., and that the differences between these states varies as a function of age and skill development. While some network dimensions may be more consequential for general maturation (and thus, are likely to correlate strongly with age), others are likely more impactful for shaping task-specific development (for example, those we observed here to be related to regulation ability). Our data preclude us from verifying this possibility, yet future work equipped with more data from a variety of different sources could successfully test this.

Our results highlight how studying whole-brain networks may inform contemporary neuroscientific models of emotion regulation. Current accounts of emotion regulation posit that individual differences in emotion regulation ability are driven by activity among a set of brain regions that comprise a dedicated network for a given regulatory strategy (Ochsner et al., 2012), while other research has only recently begun to consider the role of canonical whole-brain networks as meaningful biological units (Zhang et al., 2019). Although the notion of dedicated networks for emotion regulation strategy is enticing, our research fails to find support that a cognitive reappraisal network is associated with individual differences in emotion regulation ability. This may be because the network itself does not constitute a meaningful biological entity, despite prior evidence indicating otherwise, or because variability in both network activity and regulation ability were truncated from an otherwise full range in our sample (i.e., we only sampled a limited range of the population values of network activity metrics). Yet another possibility is that CRN activity during reappraisal may actually emerge as an extension or special form of another network (e.g., the Control Network) whose core activity might be more tightly linked with regulatory ability.

### Limitations and Future Directions

This study is accompanied by several limitations. First, although our sample size is fairly large for task-based pediatric neuroimaging, and is twice the median cell size of human fMRI studies (Poldrack et al., 2017), it is nevertheless somewhat small in comparison to other network neuroscience studies (e.g., Baum et al., 2017; DuPre & Spreng, 2017). Further, although cross-sectional research is an adequate starting point for characterizing the neurodevelopment of emotion regulation, future research would benefit from longitudinal data to examine how the trajectory of each network’s activity change across time and relate to changes in emotion regulation ability. Additionally, amid growing concerns about the reliability of computerized tasks intended to measure psychological processes and individual differences (Elliott et al., 2019; Enkavi et al., 2019), we note here that the behavioral and neural test-retest reliability of the emotion regulation task used here has not yet been established. A final future direction involves testing whether the predictive properties of the networks studied here are relevant for other emotional states, or even across different types of emotion regulation (e.g., extinction learning; expressive suppression). Addressing these concerns under further empirical scrutiny will help lend further confidence in the take-away conclusions raised by our results.

In sum, this study is the first to our knowledge that examines how the developing brain supports the acquisition of emotion regulation, a central feature of lifespan mental health. These results are the first show how canonical whole-brain networks comprise the functional architecture of emotion regulation skill acquisition across youth.

## Acknowledgments

We thank the members of the Harvard Stress & Development lab for their help in data collection. This research was supported by Grant MH103291 from the National Institute of Mental Health award to KAM. Preparation of the article was supported by a National Science Foundation Graduate Research Fellowship (NSF-GRF) to JFGM (Fellow ID: 2016220797). The authors declare no competing financial interests.

